# A Simplified Model of Motor Control

**DOI:** 10.1101/2022.11.25.517924

**Authors:** K. Arora, S. Chakrabarty

## Abstract

In general, control of movement is considered to be either cortical, spinal, or purely biomechanical and is studied separately at these levels. To achieve this separation when studying a particular level, variations that may be introduced by the other levels are generally either ignored or restricted. This restriction misrepresents the way movements occur in realistic scenarios and limits the ability to model movements in a biologically inspired manner. In this work, we propose a heuristic model for motor control that conceptually and mathematically accounts for the entire motor process, from target to endpoint. It simulates human arm motion and is able to represent functionally different motion properties by flexibly choosing more or less complex motion paths without built-in optimization or joint constraints. With a novel implementation of hierarchical control, this model successfully overcomes the problem of degrees of freedom in robotics. It can serve as a template for neurocomputational work that currently uses control architectures that do not mirror the human motor control process. The model itself also suggests a maximum threshold for delays in position feedback for effective movement, and that the primary role of position feedback in movement is to overcome the effects of environmental perturbations at the joint level. These findings can inform future efforts to develop biologically inspired motor control techniques for prosthetic devices.

## 1 Introduction

### The Complexity of Motor Control

Movement control in humans is complex and involves-numerous pathways and cellular components. Research in human motor control has navigated this complexity by developing separate branches of research that focus on specific aspects of the system in detail. For example, gait analysis [1–3] or muscle biomechanics [4–6] describe movements in terms of the trajectories taken by joints or the forces produced by muscles. These domains describe the end products of the control process, but generally donot elborate what the sources of these products are or how they shape the activity of the latter. Atthe other end of the spectrum, there are some areas that focus entirely on the nature of those sources, and describe more abstract concepts, such as how internal models of the body and environment shape movements [7–9], or how activity at the level of the brain reflects intended movements [10, 11]. These studies, in turn, do not focus on how this information is modified by spinal components or muscle properties when translated into motion. There is also interest in the role of muscle coordinations in the intermediate stage itself [12–14], but, similar to the other work, this interest rarely focuses on connections with the other areas.

### Drawbacks of Motor System Fragmentation

The bottom-up approach described above has led to important insights into how control is achieved within each of these sections, and has informed our current conceptualization of motor control. However, considering the components in isolation has limitations that constrain our understanding of motor control. To implement such a separation in each of these domains, interactions with other areas are either restricted or ignored [13]. There are multiple pathways within the motor control system to achieve the end goal of producing a movement. This redundancy suggests that a fragmented approach may miss the global features of control. An atomistic, component-oriented approach towards this system may obscure whether interactions between levels lead to different functions in the overall context [15–18].

This fragmentation has been favourable for motor control research in part because a holistic experimental investigation into the human motor control system presents considerable methodological challenges. First, without the limitations in studies that focus on a single component, it is experimentally difficult to perturb a specifically desired proprioceptive channel and to attribute observed outcomes to a corresponding element [19, 20]. Second, most tools used to record activity for motor control studies are specific to the level being investigated (eg. EEG/fMRI, EMG, video kinematic analysis). Simultaneous recording at different levels may be an option in animal models [21, 22], but its invasiveness prevents its application in human studies, and its transferability to human control principles is not guaranteed.

### The Role of Modelling Approaches

The experimental limitations mentioned here are obstacles that can be overcome by modelling. As such, models are used for applications that benefit from a conceptual representation of motor control processes, such as neuroprosthetic devices [19, 23–25]. These aim to computationally solve problems that are solved seemingly effortlessly by the human motor control system, such as the redundancy in degrees-of-freedom (DoFs). The motor system controls multiple DoFs despite existing in a three-dimensional space, a problem that has long existed in the field of robotics and requires workarounds [26]. However, facets of motor control like this redundancy (and noise, feedback interactions, etc.) span across different fragments, or levels, of motor control. Instead of the bottom-up approach that currently underlies such models, constraining them to focus on the role of a particular level, such facets may be better understood through a top-down study of movements. An improved understanding of these facets would be directly applicable to the development of prosthetic devices designed to mimic natural human movements.

### A Top-Down Approach

These ideas suggest that a more holistic, top-down investigation of the motor control system - one that includes the brain, the spinal cord, muscle activity, and the different proprioceptive feedback loops that exist at these levels - may lead to novel insights about each of their roles. Within such a model, the exact configurations of all neural circuitry along with the intricacies of their complex interactions may be difficult to replicate [27]. However, this is not necessary for a comprehensive understanding of the system as long as the underlying function of the components and interactions is preserved. As described by Marr in his three levels of understanding of complex systems [28], if such a simplified version captures the way the system responds to the overall computational problem it is trying to solve, then it may provide more functional insight than a version which accounts for each individual detail of a component’s circuitry [28, 29].

Consistent with this notion of a holistic approach, this study develops a novel, top-down model of motor control that replicates the movements of the human arm. The goal of this setup is to identify what roles these components have in controlling motion when working together. The model also aims to find out when positional feedback is useful across these levels, and which type of noise can be best overcome by this feedback. Overall, this study aims to determine whether a model built using this top-down approach can reflect what we already know about human movement while also providing novel answers to these questions.

## 2 Methods

### 2.1 Model Construction

The proposed model simulates the movements of a human arm with three joints (shoulder, elbow, wrist) that are constrained to a two-dimensional space. Given an initial configuration and a two-coordinate desired final endpoint, it iteratively computes a trajectory towards this endpoint in terms of all three angles until a specified duration. This trajectory is computed by a system of behavioural, muscle-level, and coordinate representing units (Fig 2.1).

**Fig 2.1:**
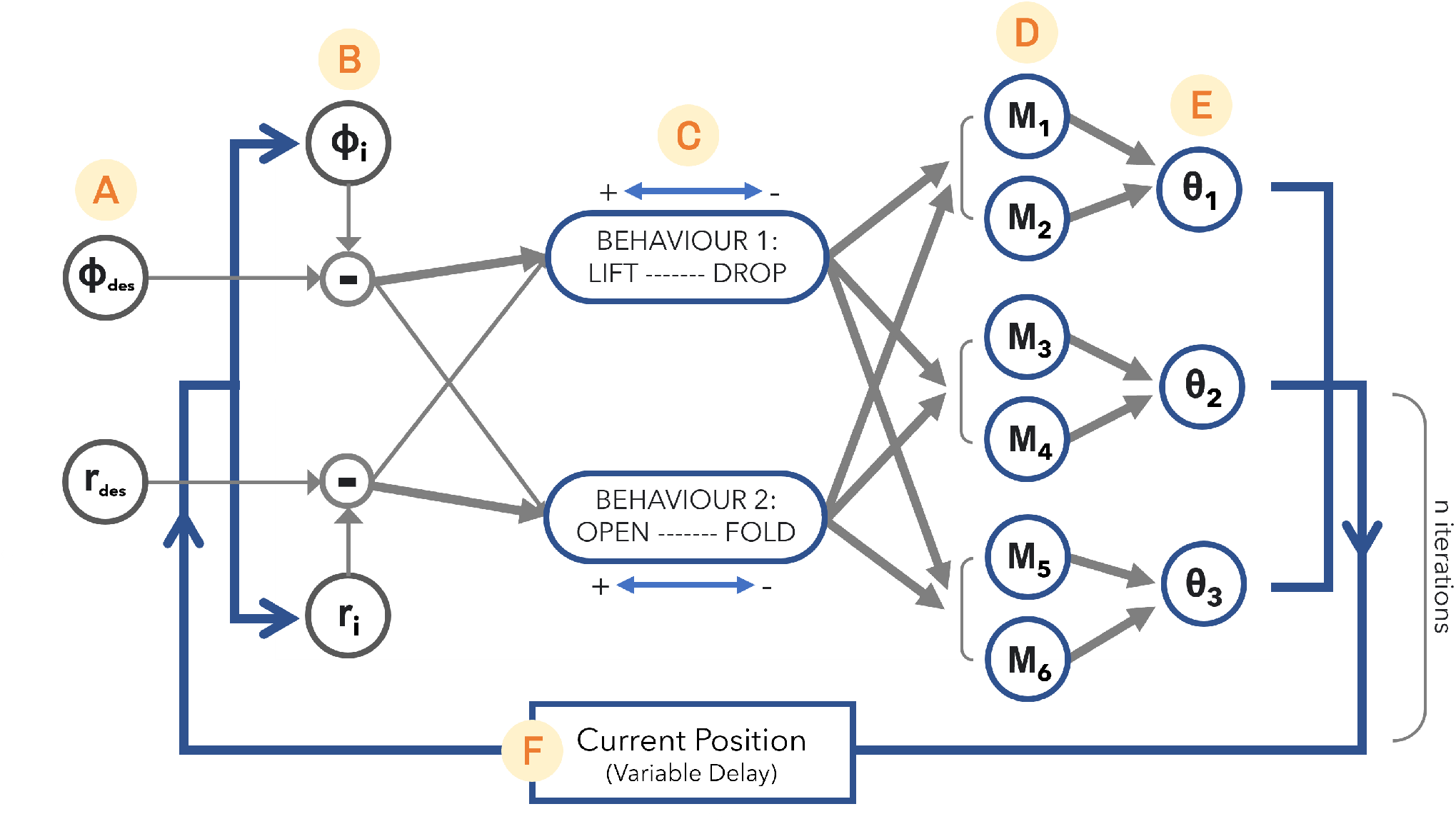
Model Schematic **A**. Desired Endpoint Coordinates **B**. Current Endpoint Coordinates **C**. Behavioural Pattern Units **D**. Muscle-level Units **E**. Joint-angle Units **F**. Positional Feedback

#### Coordinates

The desired endpoint (Fig 2.1A) and current configuration (Fig 2.1B) are used to compute the current positional disparity in terms of polar coordinates centered at the shoulder/origin (Δ*ϕ* and Δ*r*). This difference is updated at each iteration (see section 2.2).

#### Behavioural Units

This layer (Fig 2.1C) translates spatial differences between current and desired positions into muscle inputs. The activity of this layer during a movement represents the combination (and proportion) of simple behavioural patterns required to achieve any desired endpoint from the current position.

The two units in the behavioural layer represent specific, simple patterns of arm joint configurations. When active, they recruit the muscle-level units to execute movements towards their pre-defined configurations. The unit names, LIFT-DROP (ld) and OPEN-FOLD (of), reflect the opposite configurations that they propel the arm towards when positively and negatively activated respectively (Fig 2.2a-b). The unit activations are computed from the spatial differences as follows (See Table S.1 for coefficient values):

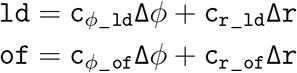

#### Muscle Units

The six muscle-level units (Fig 2.1D) occur in pairs that mimic simple agonist-antagonist muscles. The sum activation of the pair in response to a behavioural unit is constant. The proportion between the two is determined by the input from the behavioural unit, which first passes through a sigmoid function *S* (Fig 2.2c). The final muscle unit activity is determined by adding this input across both behavioural units after the sigmoid function is applied as follows:

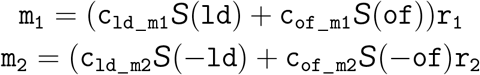

**Fig 2.2:**
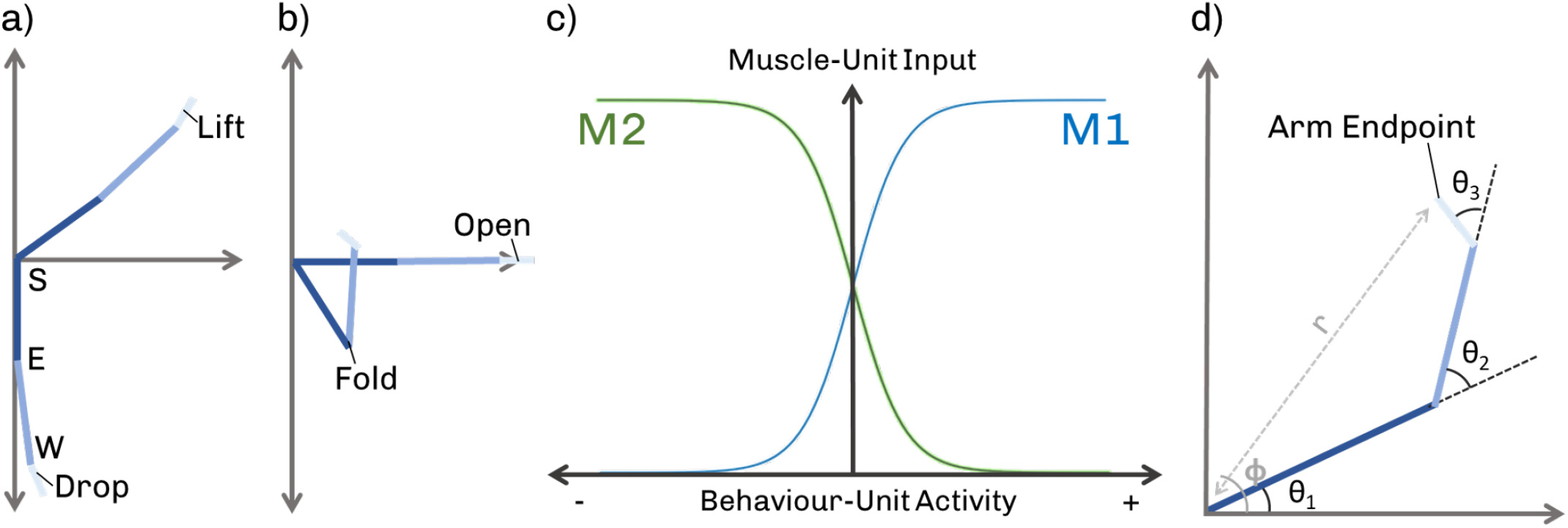
Pre-defined configurations of the **S**houlder, **E**lbow, and **W**rist **a)** for the LIFT-DROP unit **b)** the OPEN-FOLD unit. **c)** Sigmoid function relating instantaneous behavioural unit activity to a muscle-unit pair; the sum total of a muscle-unit pair’s response to a behavioural unit is constant. **d)** Endpoint co-ordinates r and *ϕ* used for positional feedback.

Similar equations are set up for all six muscles, with the relationship between a behaviour and muscle unit pair determining the sigmoid function (positive or negative) used for each unit. Each of the two units acts on one joint with equal weightage, and with the opposite effect on angular direction. The drive from previous units is scaled by a recruitment factor *r*_*n*_ to ensure smooth, continuous activation of the muscle. This factor is iteratively computed as follows:

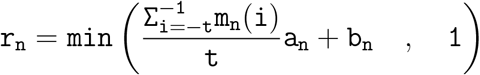

Here, *t* denotes the duration of past activity that affects the current drive. *a*_*n*_ and *b*_*n*_ control the extent of this effect. The scale factor is capped at 1, and thus never amplifies the incoming drive. It only presents the physical restriction of gradual as opposed to immediate recruitment. Lastly, signals from higher-level units were delayed to the muscles acting on the shoulder, elbow, and wrist joints by 10ms, 17ms, and 22ms respectively to reflect activation latency [30].

#### Joint Angles

Increments to joints (Fig 2.1E) are computed based on the two muscles acting on the angles (th1-3) as follows:

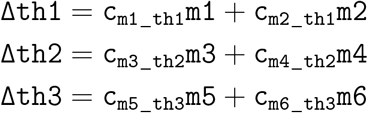

The segments (upper arm, lower arm, hand) followed a 1:1:0.2 length ratio. These lengths (in arbitrary units) are used to define the arm coordinates during analysis.

### 2.2 Additional Features

#### Feedback

Positional feedback (Fig 2.1F) was implemented to inform the model of the endpoint’s new position at each iteration. The polar endpoint coordinates *r* and *ϕ* (Fig 2.2d) were used for this purpose. Versions of the model with and without feedback, as well as with different delays in feedback arrival were considered. Without feed-back, the model does not make iterative progress; a final position is computed and trajectory to it is interpolated. Feedback delays investigated ranged from 0-15% the duration of the movement.

#### Noise

Noise was added to behavioural, muscle-level, and joint units. To the first two, signal-dependent Gaussian noise was added. To the joints, signal-independent Gaussian noise was added to mimic external sources of environmental perturbations. Versions of the model with noise being on/off at different levels were looked at.

### 2.3 Endpoint Space

Movements from various initial configurations to all points within a selected workspace were considered. This workspace is defined with respect to arm segment lengths (1, 1, 0.2 units) as the space formed by [-0.4, 1.6] along the horizontal and [-1, 1] along the vertical. Within this frame, the shoulder sits at the origin (0, 0).

### 2.4 Human Data

Model outcomes under different noise conditions were compared to human endpoint data from a bicep-curl task. Endpoints of 60 elbow flexion movements were identified from video data of a participant performing bicep curls. The mean position of the shoulder across curls was used as the coordinate origin. As input to the model, the mean endpoint achieved across these 60 points was used as a desired endpoint. All data was collected in accordance with approval by the University of Leeds’ ethics committee (ID: BIOSCI 21-007) as per the Declaration of Helsinki.

### 2.5 Distribution Comparisons

To compare the outcomes of different model parameters (different noise settings/feedback delays), and to compare model outcomes to human data, distributions of achieved endpoints (for a given initial configuration and desired endpoint) were used. The 2-D endpoint distributions were compared through the *ϕ* and *r* coordinates of their points. Estimation statistics [30] were used to identify statistical differences. In all cases, 5000 bootstrap samples were taken. Confidence intervals were bias-corrected and accelerated.

## 3 Results

The model’s movements were inspected with various initial configurations and desired endpoints across the workspace. From the movement trajectories taken by both (feedback and non-feedback) versions of the model emerged two types of endpoints: direct endpoints, that were approached correctly even without feedback (Fig 3.1a), and indirect endpoints, that were not approached correctly without feedback (Fig 3.1b). Indirect endpoints tended to be radially towards or away from the shoulder with respect to the initial endpoint. Direct endpoints tended to be those that required contributions mainly from one joint to approach, such as a simple elbow flexion. Additionally, although the arm approached direct endpoints from the correct direction in both versions, only the presence of feedback fine-tuned the arm’s position towards the end of its movement.

**Fig 3.1:**
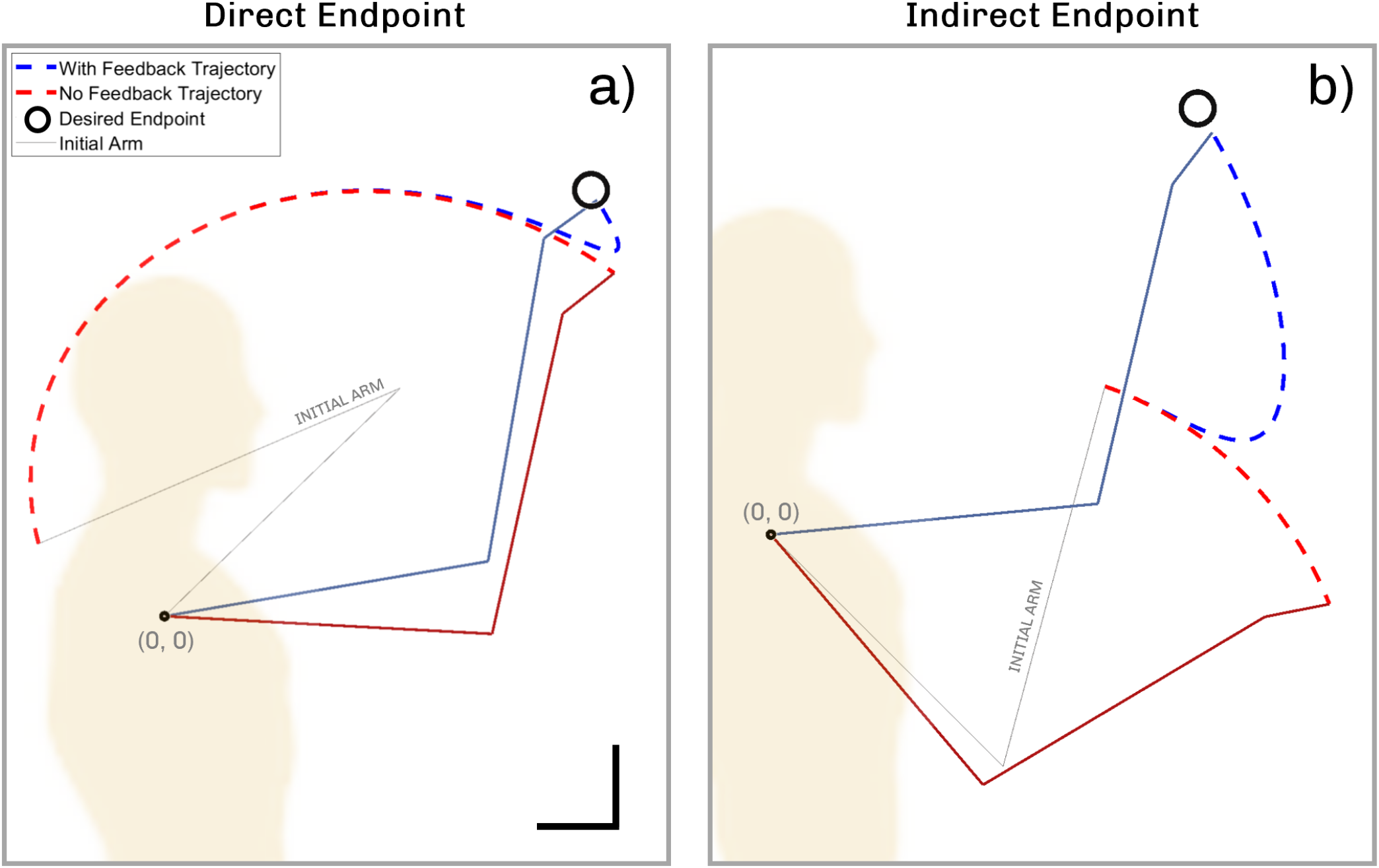
Two simulated arm trajectories with and without positional feedback. Initial and desired end-points corresponding to **a)** a direct endpoint **b)** an indirect endpoint. Scale bar 0.25 units.

These movement features were then inspected across the whole workspace (Fig 3.2) to further visualise spatial differences between direct and indirect endpoints. The model’s reaching performance shows an extension of what was described above. It was seen that the model is extremely limited in the regions which it can accurately move to without any positional feedback (Fig 3.2a-right). These limited regions, described earlier as direct endpoints, form a curved region of increased accuracy compared to the rest of the workspace. However, in the presence of feedback (Fig 3.2a-left), the entire workspace is within accurate reach.

The distance/displacement ratio was used as a measure of how linear (close to 1) or arced (higher than 1) the path taken by the arm was. It was seen that the trajectories taken by the version without feedback always maintained a distance/displacement ratio close to 1, irrespective of where the desired endpoint was (Fig 3.2b-right). In contrast, this ratio for trajectories followed with feedback was variable, depending on the desired endpoint (Fig 3.2b-left).

**Fig 3.2:**
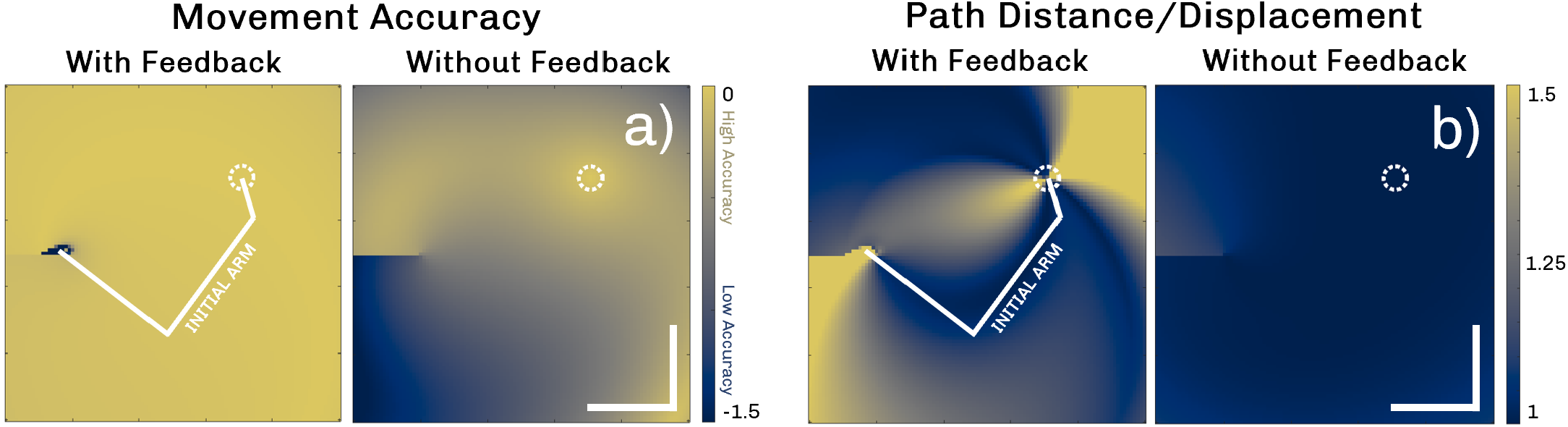
Model performance across the workspace. Heatmaps represent movements from the specified initial configuration to the respective endpoint in space. Colour values represent **a)** how close the model is able to get to the respective coordinate **b)** the distance/displacement ratio of the final path taken to that coordinate. Scale bar 0.5 units. Feedback allows accurate reaching of the entire workspace, implementing less or more complex paths flexibly as required.

**Fig 3.3:**
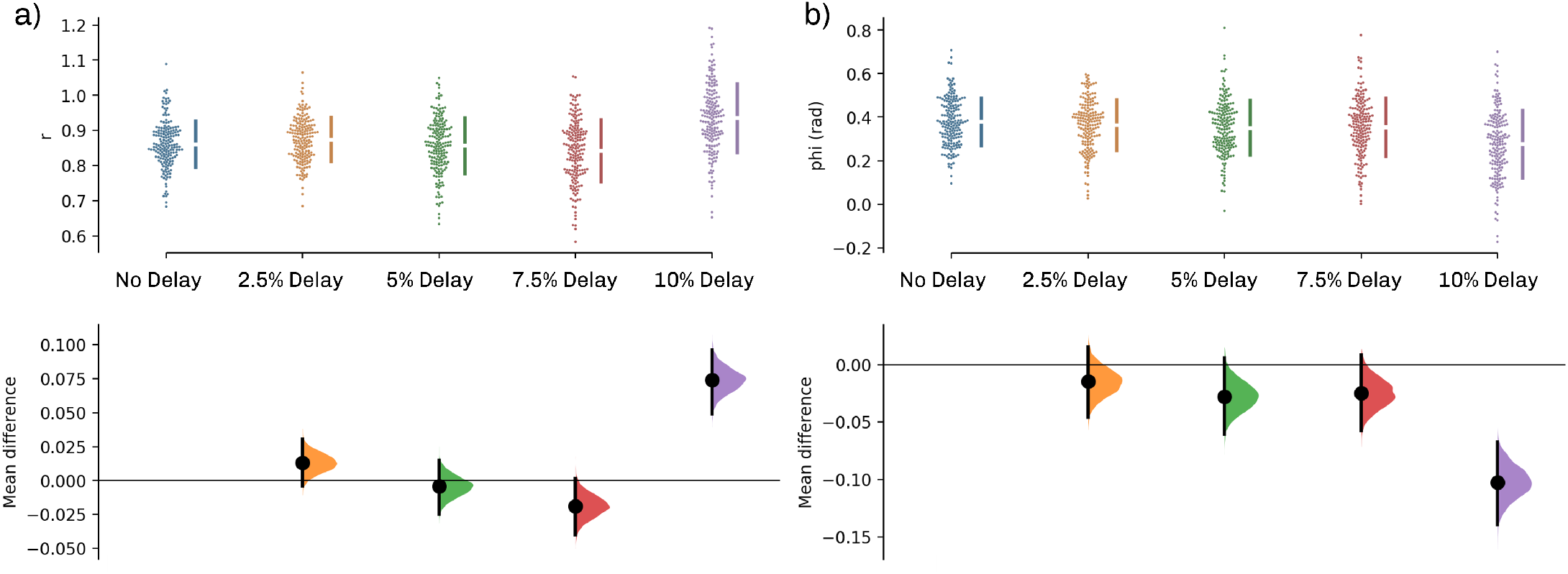
Effect of positional feedback delays: Gardner-Altman plot of differences in endpoint distributions across **a)** r and **b)** *ϕ* components with increasing delays in positional feedback. Delay times are described as a fraction of the full movement duration. **Top:** Raw data, distribution of endpoints at various delays. **Bottom:** Mean differences are plotted as bootstrap sampling distributions. Vertical error bars indicate 99% confidence intervals. Positional feedback delays do not affect model performance until 7.5%, and a threshold arises between 7.5-10% after which performance is affected.

**Fig 3.4:**
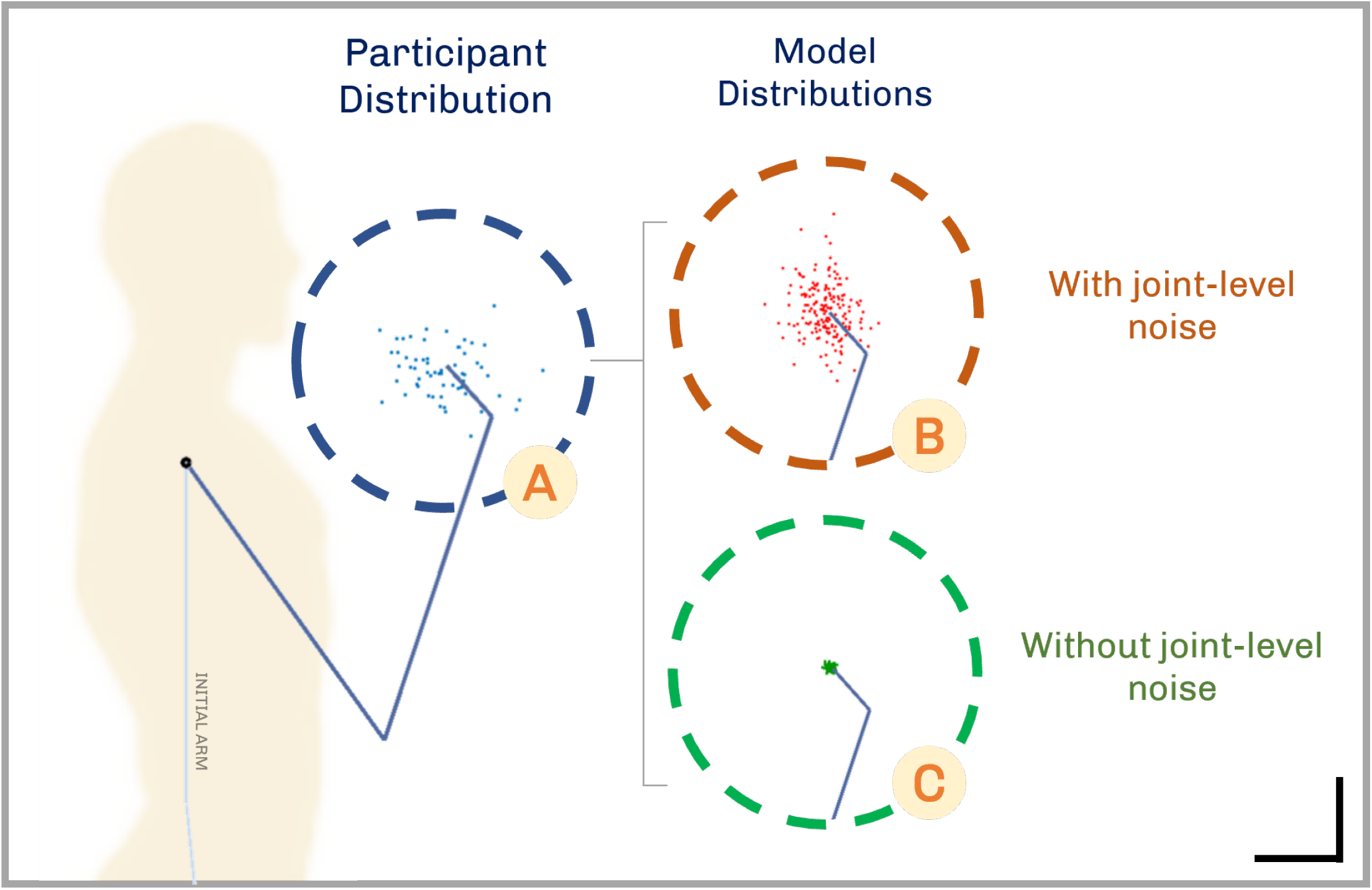
Comparison to human movement distribution. Achieved endpoint distributions for **A**. bicep curls of a human participant **B**. modelled movements with joint-level noise **C**. modelled movements without joint-level noise. All dashed circles correspond to the same region in the coordinate plane. Scale bar 0.25 units. The presence of joint-level noise is what preserves the distribution of endpoints seen in human movements.

There was no discernable pattern of gradually increasing mean differences from 0-7.5% (max. mean difference of *r* = -0.019 [99% CI -0.040, 0.013], max. mean difference of *ϕ* = 0.028 [99% CI -0.061, 0.005]), however, the distribution resulting from 10% delay is significantly different (mean diff. of *r* = 0.074 [99% CI 0.049, 0.096], mean diff. of *ϕ* = -0.102 [99% CI -0.139, -0.067]). Visual inspection of the raw data distributions showed no apparent difference in the shapes of the distributions, with each showing a shape akin to a normal distribution.

Lastly, the endpoint distributions were affected by the presence or absence of noise at specific levels. Distributions resulting from the removal of signal-independent noise at the joint level did not reflect the shape of those from human data (Fig 3.4). Without joint level noise, endpoint distributions with extremely low variance were seen. In contrast, the presence or absence of signal-dependent noise at the behavioural or muscle level units did not produce discernible changes in the endpoint distribution shape. It is noteworthy that across all noise conditions, the mean position of the distribution did not deviate from that of the participant (max. mean difference of *r* = 0.014 [99% CI -0.028, 0.052], max. mean difference of *ϕ* = 0.028 [99% CI - 0.028, 0.084]).

## 4 Discussion

In this study a top-down approach was used to develop a model of motor control that simulates the movements of a human arm with three joints (shoulder, elbow, wrist) in two dimensions. The role of positional feedback, the ability of the model to overcome delays in feedback, and the effects of noise within the model were investigated. Positional feedback allowed the arm to accurately approach all points within its reach range; without this feedback, only a limited range was accessible. Furthermore, this feedback only worked for under a 10% time delay (accounted for as a fraction of the total movement duration). Finally, it was shown that noise at the joint level was necessary to reproduce movement variations similar to those in humans.

### Positional Feedback

Model control was studied both with and without positional feedback. Without this feedback, the model failed to perform a variety of movements. The presence of positional feedback (without a temporal delay) allows the arm to precisely move towards any endpoint across the 2-D workspace. While it has been established that this is also the case for voluntary movements in humans [31], it is interesting to look at the types of movements that can still be achieved without feedback (Fig 3.1a, Fig 3.2a-right). The model suggests that for basic movements (which mainly require only one joint, or only a change in *ϕ* from the original position), positional feed-back might not be an absolute requirement for completion; such endpoints are approached correctly by the model even without feedback.

Given that these movements are completed by the model in less time compared to more indirect ones, they can be considered to represent ballistic movements [32]. These ballistic movement properties are well represented in terms of how the model’s movement without feedback follows a more direct, undeviating path (low distance/displacement ratio) rather than displaying arcs with increased curvature [33]. This comparison could later be examined by quantifying velocity and acceleration patterns of the movement endpoints and segments over time.

### Feedback Delays

We have presented the results here showing evidence of precision being dependent on time delays in positional feedback. Addition of a feedback delay did not gradually decrease model precision; rather, a delay threshold arose up to which performance is maintained at a uniform level and beyond which endpoints cannot be successfully achieved.

The fact that such a threshold exists provides a conciliatory explanation for conflicting work that used short time delays and suggested that feedback delays do not affect endpoint behaviour [34], and work which has suggested that they do [35].

However, the value of this threshold suggested by the model is not consistent with such previous reports. This delay threshold is between 7.5%-10% of the duration of the full movement (Fig 3.3). Assuming a voluntary arm movement takes between 200 - 300 ms depending on the situation, the proposed maximum functional feed-back is in the order of 20-30 milliseconds. Depending on the duration of the task, this may be shorter than it actually takes for positional information to travel the full route to the brain [35, 36]. It has been suggested that for sensory feedback to effectively overcome time delays, its coupling with internal models of movements and the environment is necessary [7, 37]. It is likely that the model is more vulnerable to delay-caused deviations in accuracy because such internal representations were not implemented here. However, it is also interesting to note that the maximal delay threshold suggested here does compare well to positional information travelling to the spinal cord, as per short-latency reflex onset times [38, 39].

This instantaneous change across feedback delays confirms that the results described above would also hold in a more realistic system where this delay is present (and below threshold).

### Environmental Perturbations

To replicate environmental perturbations, different feedback and delay conditions were compared using both signal-dependent noise within the system, and signal-independent noise (environmental perturbations) at the joint level. This was validated by comparison with human data (Fig 3.4).

In the absence of environmental perturbations at the joint level, the model maintains sharp precision across noisy repetitions. In this case, the distribution of end-points no longer reflects the distribution of the collected human movements. This implies that positional feed-back accounts for these perturbations rather than for signal-dependent noise within the system. The data used here involved bicep curl movements with a 4 kg weight at a set speed. The presence of a weight, the fixed time frame of movements, and the fatigue induced by performing 60 bicep curls, are all factors that could potentially affect how natural these movements are in comparison to what this model is attempting to replicate. In the future it may be more suitable to create a dataset tailored for comparison to this model.

### Hierarchical Control

The effect of different feedback, noise, and delay conditions has already been described. The form of hierarchy used to control a higher dimensional joint-space as opposed to the coordinate workspace is the main feature at the core of all these versions of the model.. Though the notion of hierarchical control has been used to create successful models of control in the past [40], most have conventionally implemented biologically-inspired components at either the brain [41] or muscle synergy [13] level alone. Second, regardless of the biological focus, these models are more complex than the one proposed here. This model investigated whether it would be possible to replicate human arm movements using simple representation of motor control components: the highest level only as current/desired coordinates and the difference between them, the spinal level only as a pair of pre-known arm configurations that recruit a higher number of muscles (similar to muscle synergies), and muscles only as agonist-antagonist pairs that act on one joint.

These characteristics distinguish the proposed model from works with similar control objectives, and the observations show that this system with a modular format performs well. This model validates a novel control strategy for high-DOF movements in lower dimensional spaces.

### Relationships to Current Work and Future Applications

The proposed model provides insights into functional interactions between different biological components, while also serving as a tool to replicate control. Its modular nature holds even more potential for future biologically-driven motor control modelling.

In spite of its current simplicity, the model captures dynamic features of human movements. For example, it demonstrates functional differences between ballistic and fine-tuned movement, based solely on the location of the desired endpoint with no additional information. It also reproduces oscillatory tremors during reaching as a function of positional feedback delays (See Fig S.1).

Currently, the model covers cortical and cerebellar components via coordinate representations, but not regions concerned with intention. Goals and action selection add a dimension of control that a coordinate alone does not provide; a coordinate may need to be reached for (simple arm movement modelled here) or walked towards (repetition requiring a different approach). Goals, intentions, and action selection have been extensively studied at the higher levels [7, 41], but their models tend to be limited in their connectivity and interactions with downstream control. The model presented here is unique in that it can be coupled with adapted models of internal goals and plan selection. This would produce a model that can explain and replicate not only the physical act of control, but also how internal choices affect these biological signals and how different goals are executed.

Furthermore, the current model is static. None of the weights used change situationally or over time. In order to replicate features such as learning in novel tasks, or fatigue in muscles through extended use, temporal changes should be included. A future version of the model may use modelled intention as described above to dynamically change the system’s predisposition to specific behavioural patterns. It may also show changes in favoured muscles if those in use develop fatigue, assuming a redundancy in the muscle layer. This line of investigation would use the vast set of available artificial neural network techniques, used extensively in movement control [42–45]. These are a good computational fit for systems with layer-organised units, use iterative feedback, and may change connectivity based on the progressive fulfillment of a specific goal. All of these criteria are met by the motor control system as envisioned by this model. Converting this static model into a dynamic neural network would bring it closer to a tool directly applicable to devices that control and execute movements.

Finally, a biomechanical connection, i.e., measures of forces and joint torques produced during movements, could make the model more robust. This could be combined with previous research on movements and muscle efficiency [46] to develop optimization criteria for the computational tools described above.

An evolved version of the top-down approach may include all the above-described features. This would simulate biologically-informed movements at all levels of the motor control system, from goal-selection and planning to muscles and joints. Such a framework could serve as a universal template for a variety of motor control modelling applications. Novel neural mechanisms of dysfunctional gait or movement tremors could have their effects on joint dynamics or reaching patterns tested with relative ease. Variable signal delays could provide evidence for whether certain regions of the brain or spinal cord play a greater role in executing particular movements. Importantly, the modularity of this model allows it to benefit applications that require more than its basic form. Complex decisionmaking models and learning could be simply linked to this template to provide a clear understanding of their effects on the peripheral systems. The applications discussed here are immediately applicable to research streams focused on differentiating motor dysfunction, and those that aim to use neuroprosthetics to remedy these dysfunctions.

Such universal, modular, and accessible templates hold great promise for accelerating research in a given field, be it the role AlphaFold plays in identifying protein configurations [47], MIIND in dynamic neuronal modelling [48], Open Source Brain in simulating brain regions [49], and several others. A future version of the model proposed here may offer similar opportunities for accelerating the field of motor control.

## 5 Conclusion

The goal of neurorehabilitation research has frequently been to understand and implement human control *in silico* to facilitate devices that strengthen lost motor function, with a focus on data-driven approaches as a foundation. This model offers a more biologically-informed, holistic resource for similar future work.

This model was able to successfully control a 2D arm using a novel hierarchical architecture, providing insights into the nature of movements, feedback, and noise within this system. The simple top-down approach implemented to model the arm’s use of its three joints to reach a desired endpoint produced near-perfect accuracy. It chooses a more or less complex path to carry out a naturalistic movement based solely on the relationship between its current and desired positions in space. It also fine-tunes its position as it approaches the endpoint.

Despite having very few units and no built-in optimisation techniques, the proposed model successfully mimicked known, functional features of movement. This suggests that a biologically-inspired top-down approach is a powerful way to gain insights about human movements and the roles of different levels within them. This model can possibly be scaled up to 3D movement and the entire human body by adding more behavioural units to account for the increased number of muscles and joints involved. Using this system as a foundation, future versions that expand on it and include more components of control could be developed. These could be used to link specific biological causes to phenotypes and movement characteristics. These could also be used to quantify the impact of injuries and potential treatments on control ability. Future modelling work can undoubtedly unlock more of the potential within the top-down approach, and the model proposed here serves as the foundation for such possibilities.

## Supplementary Material

All model code and data used for figures can be obtained through the following link: https://github.com/arora-k/Simplified_Model_of_Motor_Control. All model code was written in MATLAB [50].

## Appendix

**Oscillatory Tremors**

**Fig S.1:**
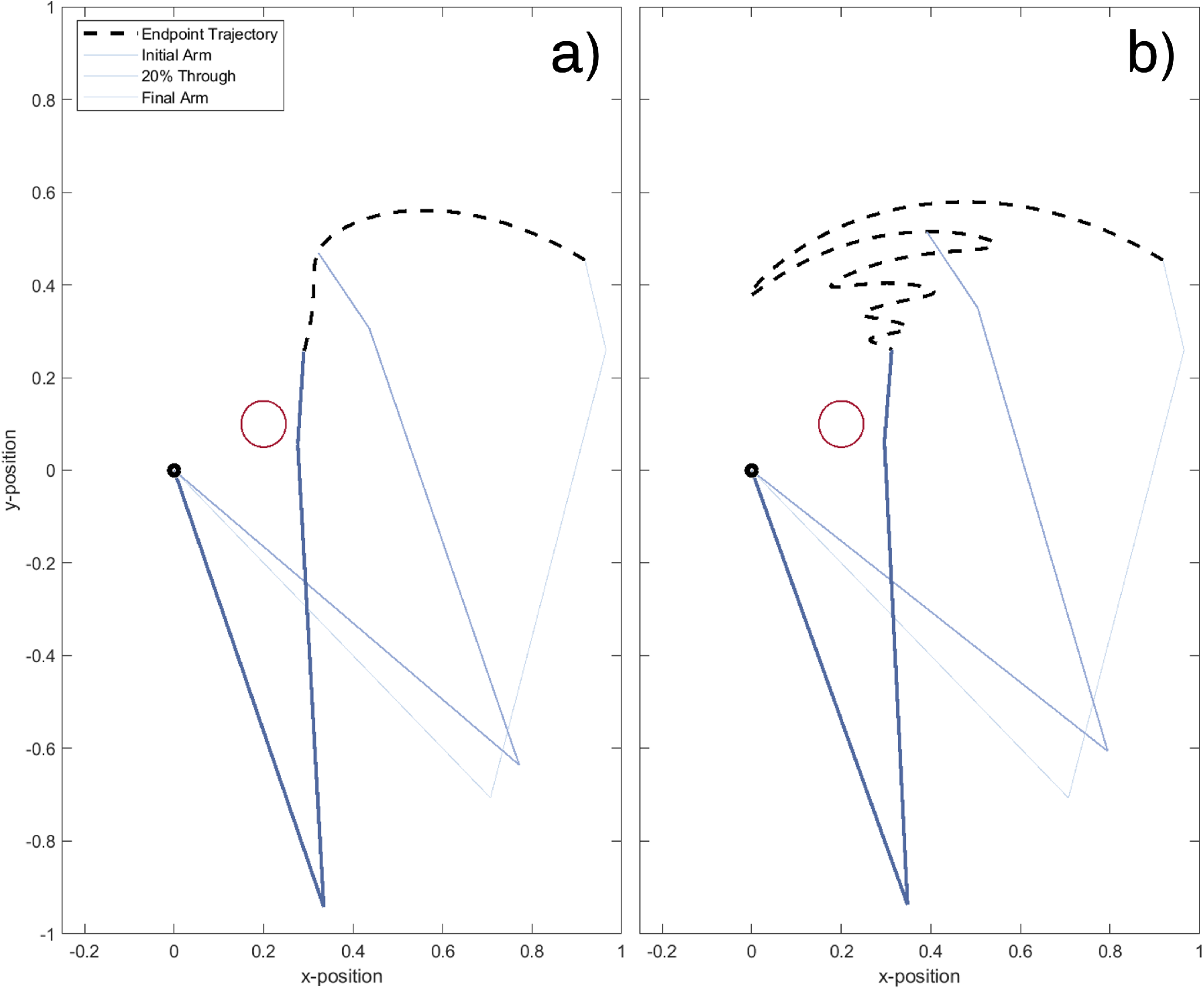
Example movement simulated by the model with positional feedback at **a)** 0% delay and **b** 6% time delay. Delayed path shows oscillations about the path taken with no delay, but the magnitude decreases over time and the desired endpoint is still approached.

**Table S.1:**
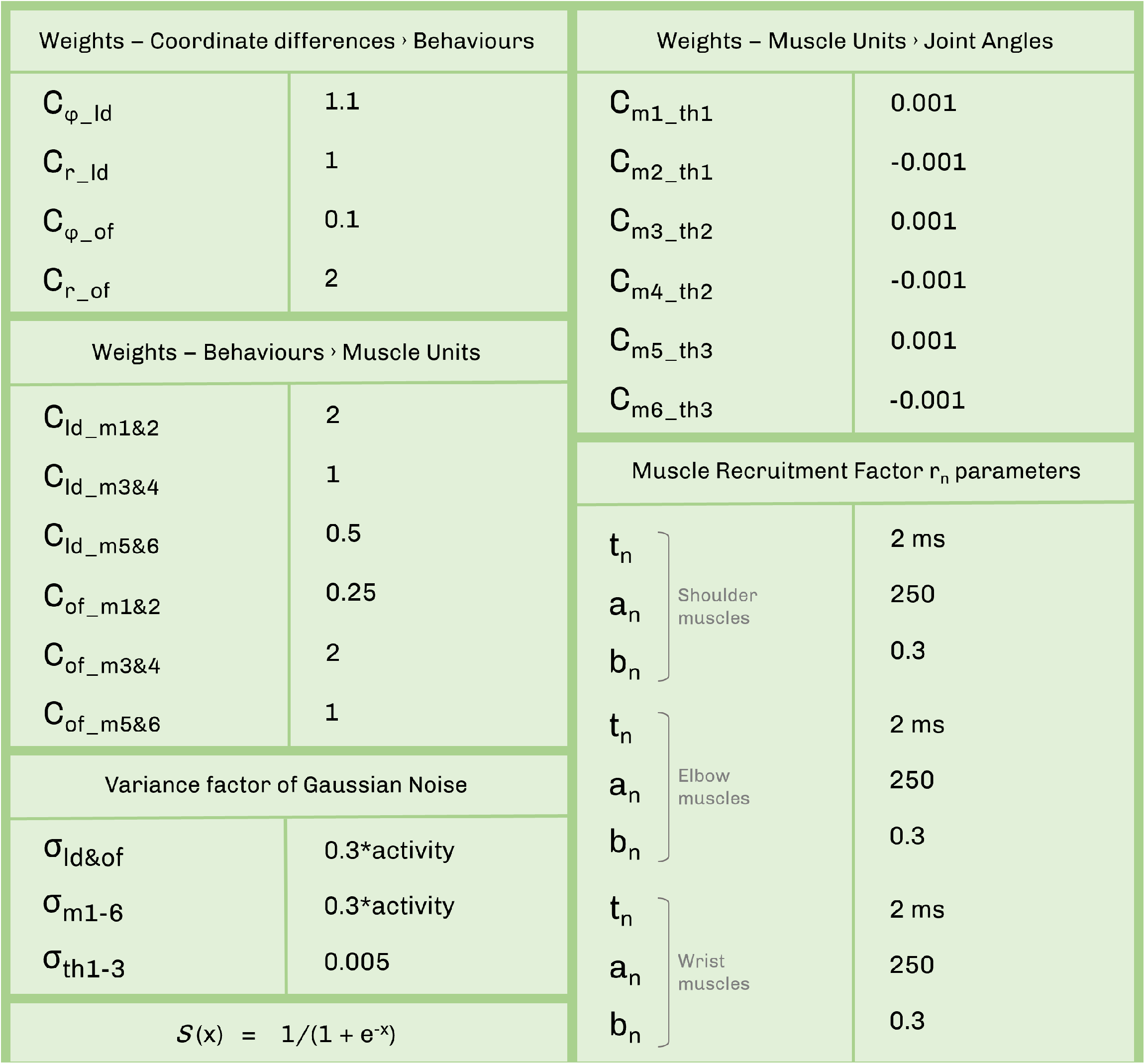
Model Weights and Constants

